# Preparation of cationized albumin nanoparticles loaded indirubin by high pressure hemogenizer

**DOI:** 10.1101/2021.05.15.444280

**Authors:** Houra Nekounam, Rassoul Dinarvand, Rahele Khademi, Roya Karimi, Hossein Arzani, Narges Mahmoodi, Elham Hasanzadeh, Morteza Kamali, Masood Khosravani

## Abstract

Indirubin can be applied as an anti-cancer drug for inhibition of brain tumors. However, its performance is reduced due to hydrophobicity. In this study, we synthesized cationic human serum albumin (CHSA) nanoparticle by a new hybrid approach for improvement the surface chemistry of albumin and investigate the amount of indirubin loaded CHSA nanoparticle. In this study, the generated mechanical force from a high-pressure homogenizer (HPH) was used to make nanoparticles with a certain size with narrow polydispersity. The results indicated that the size of indirubin loaded CHSA nanoparticles were 130 nm and their zeta potential were +9. Besides, the encapsulation efficiency and drug loading capacity were found to be 85% and 5.8 %, respectively. To the best to our knowledge, this is the first time that indirubin has been used in albumin nanoparticles. In this study, indirubin loaded CHSA nanoparticles was shown can be a potential candidate for drug delivery in the treatment of glioblastoma. Moreover, the cationized form allows the chemical agent to be transmitted to the brain.

## 1. Introduction

One of the progressive types of cancer are malignant brain tumors which lead to death in most patients approximately one year from the time of diagnosis in early stages [1, 2]. The current treatments normally including brain surgery combined with irradiation treatment with or without chemotherapy are restricted due to the tumor’s biology and its location in the brain within approximately 2 cm of the original lesion [2, 3]. Despite the recent advancements in new chemotherapeutic agents and regimens for cancer treatment in other parts of the body [4], chemotherapy of brain tumors has been poor due to the blood-brain barrier (BBB)[5].

The BBB is essential for the maintenance of the neuroparenchymal environment, however, represents a barrier to the entrance of therapeutically drugs [6]. The BBB includes various cell types such as pericytes, astrocytes, endothelial and microglial cells[6]. The tight junction proteins between the brain capillary endothelial cells are mainly responsible for static barrier properties, resulting in the poor transfer of almost all drugs [6]. Besides, ATP-binding cassette efflux transporter P-glycoprotein play a key role to transport the huge chemotherapeutic agents and hydrophobic compounds outside the brain capillary endothelial cells[7, 8]. To overcome this problem, the nanostructure can be used as colloidal drug carriers to enhance the bioavailability of drugs by increasing their diffusion through biological membranes and protect them against enzymes. Among colloidal drug carrier systems, protein-based nanoparticles have attracted great attention due to certain advantages such as higher stability, being non-antigenic and non-toxic and their easy to scale up during manufacture [8-12].

Human serum albumin (HSA), the most abundant plasma protein, is considered as a non-toxic, non-immunogenic and biodegradable protein carrier with a half-life of almost 19 days, which is soluble in water allowing ease of delivery by injection [2, 8, 13, 14]. Therefore, many metabolic compounds and therapeutic drugs can be transported by this nano-carrier due to size and abundance. It can be also considered as a nano-carrier for the administration of higher doses of a chemotherapeutic agent for cancer patients with improving anticancer efficacy without increasing side effects [13, 15]. Kim et al. [16] reported that curcumin loaded HSA nanoparticles with size distribution of 130–150 nm have higher therapeutic effects than curcumin in tumor tissues of mice without inducing toxicity. It can be attributed to the ability of albumin to cross the endothelial cells membrane. In our previous work [2], we also reported that imatinib base loaded HSA nanoparticles can be introduced as a potential candidate for drug delivery in the treatment of glioblastoma due to greater cytotoxicity effect on U87MG glioblastoma cells in comparison with lonely imatinib base.

The cellular uptake albumin can be increased by cationic modification [17]. Cationic bovine serum albumin (CBSA), a typical example of absorptive mediated transcytosis, is considered as a potential brain targeting carrier [18-20] due to interaction of the positive charge around the albumin with the negative surface of the brain capillary endothelium [21], leading to across the BBB. Cationized albumin was used for β-endorphin to cross the BBB[22, 23]. Brain uptake of β-endorphin enhanced after conjugation with cationized albumin. Researchers evaluated the siRNA delivery system to treat metastatic lung cancer [24, 25] and the modification of serum albumin with the cationic groups allows control and surface charge for optimized drug delivery. Such modification also leads to more efficient targeted delivery systems without increasing the toxicity [18, 26].

The homogenizing valve is usually used for cationization which contains a pump that conducts fluid under pressure through a small orifice between the valve [27].

In the process of drug synthesis industrially, the high-pressure homogenizer (HPH) is an important especially tool for size reduction, mixing, and stabilization of dispersions such as macro- and micro-emulsions, and suspensions. High local stresses created by HPH causes an effective decline in particle size. Another feature is its ability to scaling up due to continuous production[28, 29]. In HPH the liquid premix is passed through a narrow split under high pressure. This premix could be a coarse emulsion, micro-suspension, or dispersion[30]. The essential factor for size reduction and droplet breakup is homogenizing pressure. when the premix passes under high pressure through a small orifice [30]. During the transmission of liquids through the HPH, the collected molecules that are associated with both the liquid and solid interacting could change the molecular dynamics of liquids[30, 31]. Understanding the importance of HPH processing parameters is essential due to the impact of these changes on physicochemical properties such as size, dispersion index (PDI) and surface charge, which affects the stability and cellular uptake[32].

The special and correct connection of molecular fluids transferred through the HPH device causes the development of colloidal formulation with eligible pharmaceutical applications[30].

Indirubin, the active ingredient of the Danggui Longhui Wan, is a bis-indole compound and protein kinase inhibitor [33, 34]. The drug has different pharmacological effects, including anticancer, anti-metastatic and anti-inflammatory properties [33, 35] . In the clinical setting, it is used to treat chronic myelocytic leukemia [33, 34]. It has been also shown to have considerable antiproliferative properties and induce apoptosis in multiple tumor cell types such as cervical, liver, and lymphoma cell lines [35]. Besides, drugs of the indirubin family may improve therapeutic effect in glioblastoma owing to the inhabitation of two major hallmarks of the malignancy, tumor-cell invasion and angiogenesis, through the effects on cell migration. Therefore, indirubins can be considered as high potential for glioma treatment by affecting tumor and endothelial compartments [36]. In this work, we synthesize CHSA nanoparticle using desolvation and nanoparticle albumin bound (nab) technology methods to improve the surface chemistry of albumin. We investigate the possibility of loading indirubin in nano-carriers of albumin and surveyed factors related to the degree of encapsulation and release, how the drug was released. Since indrobin is first loaded on a nano-carrier protein we were able to obtain good results in terms of size, zeta potential, drug loading, release.

## 2. Materials and methods

Albumin was purchased from Sigma-Aldrich (USA) and indirubin from Baoji Guokang Bio-Technology Co Ltd (China). Cell lines were obtained from the National Cell bank (NCBI) of Pasteur Institute (Iran). Ethylene diamine(EDA), hydrochloric acid (HCl), sodium acetate, acetic acid glacial and 1-Ethyl-3-(3-dimethyl aminopropyl) carbodiimide (EDAC) 3-(4, 5-dimethylthiazol-2-yl)-2, 5-diphenyl tetrazolium bromide (MTT) were bought from Sigma-Aldrich (USA).

### 2-1. Preparation of indirubin-CHSA nanoparticles

CHSA was prepared according to the described method by Thole [37]. Briefly, the pH of 50 ml EDA solution (0.9 M) was adjusted to 4.75 and 800 µl HSA solution 50% (m/v) was slowly added in an ice bath and pH of final mixture set to 4.75 again. acetone solution as the solvent of indirubin and 72 mg from EDAC were added while stirring at an ice bath for 2 hours. The reaction was terminated by adding 260 µl of 4 M acetate buffer (pH 4.75) and extensively dialyzed against double deionized water and homogenized by High-Pressure Homogenizer (IKA, LABOR PILOT 2000Bar, Germany) as much as 2000 psi for nine cycles.

The obtained nanoparticles were concentrated with an AmiconR Ultra-15 centrifugal filter device. The resulting colloid was rotary evaporated to remove acetone at 25°C for 15 minutes. The solvent was removed by lyophilization for 24 hr, at -55 °C and the powder of obtained indirubin-HAS nanoparticles was stored at 4°C for the long-time storage.

### 2-2. Characterization of indirubin-CHSA nanoparticles

#### Particle size and zeta potential measurements

The mean diameter of indirubin–CHSA nanoparticles was measured via the dynamic light scattering (DLS) method (Malvern Zetasizer ZEN 3600). The samples were diluted with double deionized water at a ratio of 1 to 10 and measured at 25 °C with a scattering angle of 90°. Then the zeta potential was determined using electrophoretic laser Doppler anemometry using a zeta potential analyzer (Malvern Zetasizer ZEN 3600) and the size distribution was determined using a polydispersity index (PI). IEF analysis. CHSA was characterized according to the PI with an LKB Multiphor II isoelectric focusing (IEF) apparatus (Pharmacia, Uppsala, Sweden) which is equipped with AmpholineTM PAG plates and a pH gradient from 3.5 to 9.5 (GE Healthcare, Amersham, UK) to verity the cationation of CHSA. NaOH and H3PO4 solutions were used as cathode and anode, respectively SEM analysis. The morphology of nanoparticles was carried out by scanning electron microscopy (Hitachi S-4700 FE-SEM; Tokyo, Japan) after sputtering with gold.

#### Determination of drug encapsulation efficiency and drug loading capacity

The indirubin encapsulation efficiency was defined as the weight fraction of the indirubin loaded CHSA. To measure the encapsulation efficiency, 1mg of indirubin loaded CHSA nanoparticles was dissolved in 10 ml of ethyl acetate/propanol (9:1, v/v) and sonicated for 30 minutes for the extraction of indirubin. The amount of indirubin in the solution was measured using UV-visible spectroscopy (Optizeri 2120 UV plus). The results were compared with the standard curve and extended to 10 ml supernatant and the encapsulation efficiency (EE%) was calculated using the following equations.

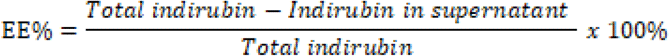

EE% is considered as the ratio of the weight of indirubin encapsulated into a CHSA to the total indirubin added whereas drug loading capacity is the ratio of the indirubin to the weight of the total CHSA.

The loading capacity (LC%) was determined as the following formula:

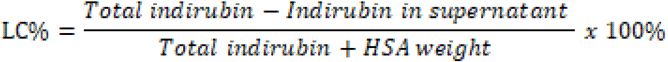

#### The drug release profile of the indirubin loaded CHSA nanoparticles

To understand the model of indirubin release from CHSA nanoparticle, 2 mg/ml indirubin-CHSA nanoparticles were solved in double deionized water. Subsequently, the prepared sample was inserted into a dialysis bag and immersed in 100 ml PBS at 37 °C (pH 7.4) for 24 hours under constant stirring with a shaking rate of 100 rpm. 2 ml of the sample was drawn out from the medium at spesific hours intervals and the same volume of fresh medium was added. Each sample was mixed with 6 ml of ethyl acetate, vortexed for 12 minutes and centrifuged at 1500 rpm for 5 min. Then, the obtained supernatant was characterized three times for each sample using UV-vis spectroscopy with the absorbance at 290 nm (Optizeri 2120 UV plus) to measure the concentration of the released drug.

## 3. Results and discussion

### 3.1. Characterization of the indirubin loaded CHSA nanoparticles using DLS, IEF, SEM, and DSC

Various methods such as coacervation/desolvation [38],emulsion-based methods [26]and nanotechnology albumin-bound [39] have been employed for drug loading into cross-linked albumin molecules or on their surface. The main method used in this synthesis is a combination of both coacervation/desolvation and nab technology methods. This is because nab technology by HPH apparatus acts as a cross-linker instead of toxic glutaraldehyde that existing in the coacervation/desolvation method.

DLS analysis demonstrated that the mean sizes of indirubin loaded CHSA nanoparticles were 130 nm. Also, the SEM image of the indirubin loaded CHSA nanoparticles shows almost uniformity shape with the range of 90 to 220 nm (Figure 1a). As shown in Figure 1b, most nanoparticles have a size of almost 130 nm which is according to DLS results. According to the literature, in the most solid tumors, nanoparticles with the size of even 300 nm can extravasate into the tumor interstitial through the hyper permeable vasculature in the most solid tumors [40] and the nanoparticles with the size of even larger than 200 nm were also used to transport drugs to the BBB [6]. Therefore, indirubin loaded CHSA nanoparticles with a size of 130 nm can be suitable for drug delivery across BBB. Researchers stated in synthesis with the HPH, the particle size of CHSA NP was impressively large (about ∼800 nm) [41]. We also saw the same result; they filtered by 200 nm pore size, to reach the average size of 198nm. (Byeon et al. 2016).

**Figure 1:**
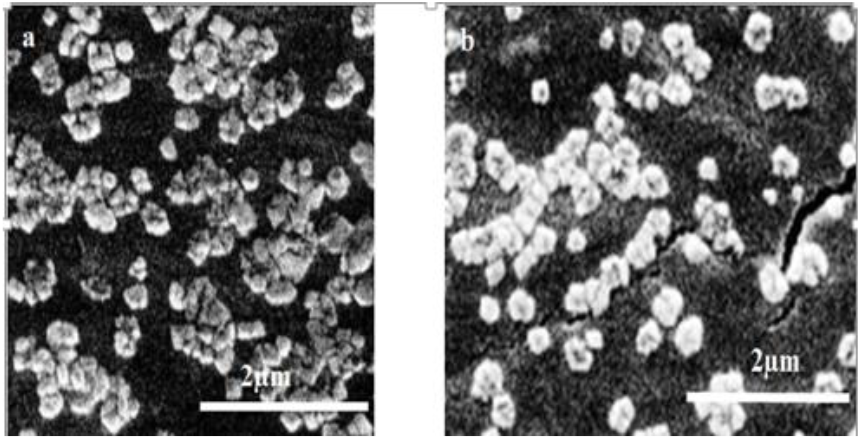
(a,b) Scanning electron microscopy (SEM) picture of indirubin loaded CHSA

**Figure 2:**
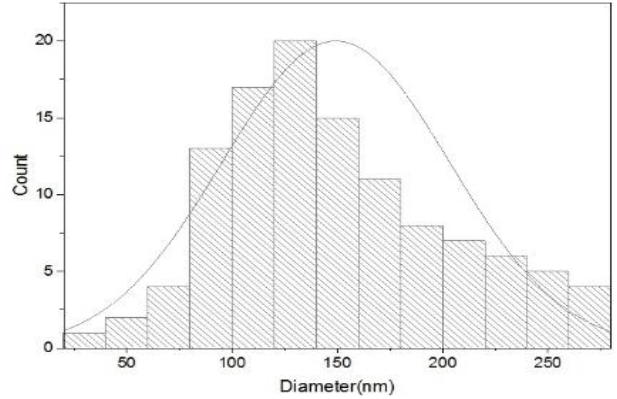
Particle size distribution histogram indicating expected particle size diameters

Moreover, the zeta potential of indirubin loaded CHSA nanoparticles increase from negative values of native HSA approximately -21.4 ± 1.3 mV to positive values +9 ± 0.5 mV which can confirm the cationized HSA due to the presence of positive charges. This value still is appropriate (within 10 mV) with the optimal zeta potential for in vivo transmission and delivery of nano-carriers [20].

Therefore, indirubin loaded CHSA can be electrostatically bound to the negatively charged residues on the BBB. The literature indicates that the cationic albumin surface undergoes absorptive mediated transcytosis of nanoparticles [42]. PI of cationized proteins was assessed by immobilized pH gradient gels in an IEF apparatus which showed a shift in the PI value of native albumin approximately 4.0 to about 9 for the cationized form Figure 3. (IEF Marker Figure 3)

**Figure 3:**
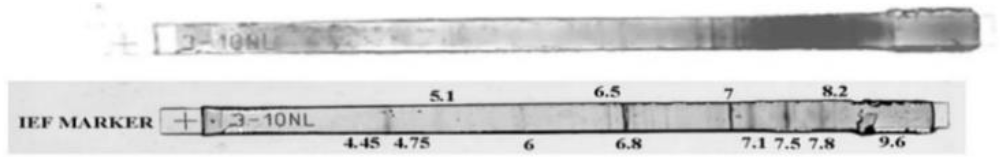
IEF strip of CHSA nanoparticles and IEF Marker.

### 3.2. Investigation of encapsulation efficiency, drug loading capacity, and release profile

EE% and LC% are considered as two important parameters in drug delivery. The EE% and LC% were 85% and 5.8%, respectively. The release profile of indirubin loaded CHSA nanoparticles were investigated in vitro at 37 °C in PBS at pH 7.4. (Figure 4). The results demonstrated that the indirubin release from CHSA nanoparticle was about 35%, 67% and 82% after 4, 10 and 18 hours, respectively which indicated a fairly slow release profile and a sustained release of indirubin from the CHSA nanoparticles. Half of the drug content was released at the first 4 hours. The release profiles were similar to imatinib loaded HSA [2] and aspirin loaded HSA nanoparticles [43]. Also, the release results were similar to CPT loaded BSA-PMMA nanoparticles which were investigated by Maofang Hua [44]. However, in this study, the initial drug release is slower for the structure of the polymer but the major of the drug in both models is released within 20 hours (Figure 5).

**Figure 5:**
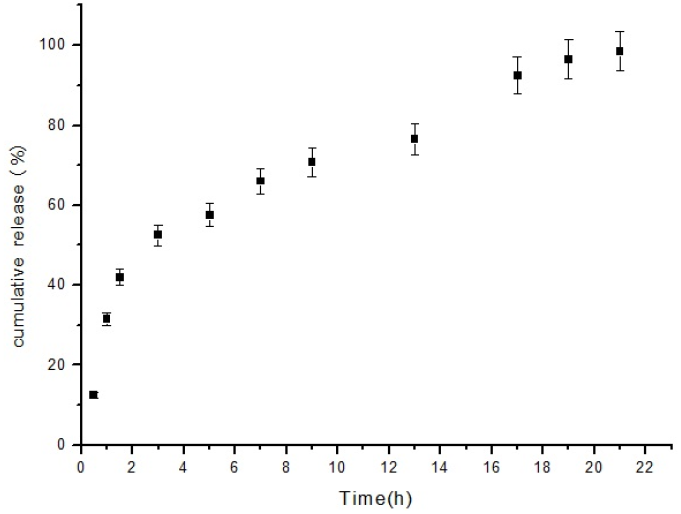
The release profile of indirubin from CHSA nanoparticle

The standard curve of DMSO/indirubin solutions is shown in Figure 6.

**Figure 6:**
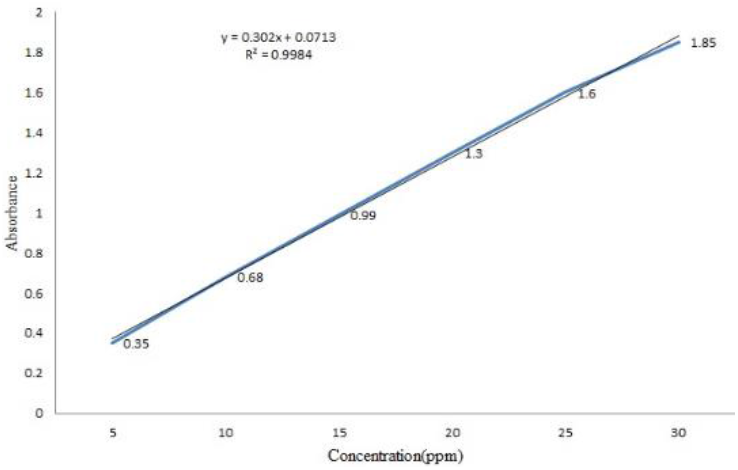
Standard curve of indirubin-DMSO Solution

### 3-3. Increase solubility of indirubin in water solution

Free indirubin is sparingly soluble in aqueous solutions. While macroscopic undissolved components of the indirubin can be seen in the solution (Figure 7a); but, indirubin-albumin nanoparticles solution is clear by light pink color. We used a higher concentration of free indirubin and no observed mild pinkish color in water (Figure 7b).

**Figure 7:**
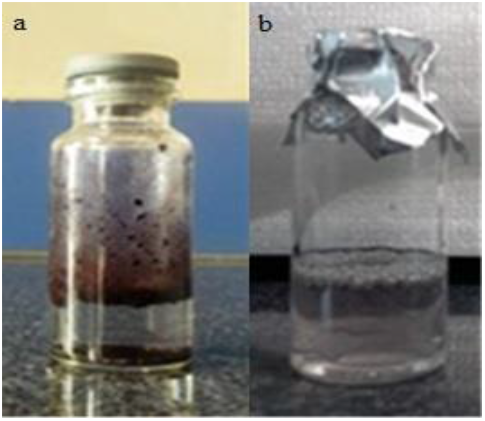
Increased solubility of the drug in an aqueous medium in nanoparticle form

## 4. Conclusions

This study investigated indirubin-CHSA NP as an effective drug for treating brain tumors. Indirubin is introduced as a useful anti-cancer, anti-metastatic and anti-inflammatory drug. However, the therapeutic effects of anti-cancer drugs decrease for the glioma treatment due to ATP-binding cassette efflux transporter P-glycoprotein. Therefore, nanoparticles are proposed owing to their tumor-targeting properties and internalization efficiencies. Our experiments exhibited that the mean sizes and the PDI of indirubin loaded CHSA nanoparticles, made with the new hybrid construction method were 130 nm (range of 90 to 220 nm) and 0.2, respectively. The use of the HPH caused uniform structure and size. This method is suitable for scaling up in industrial production. Likewise, the zeta potential of the indirubin loaded CHSA nanoparticles was positive which indicates the cationized HSA. Our results also indicated that the EE% and LC% were 75% and 5.8%, respectively. CHSA had an important role in the fairly slow release profile of the indirubin loaded CHSA nanoparticles. These nanoparticles had a considerable cytotoxic effect on glioblastoma. As well as increase the solubility of the compounds promises to improve the therapeutic effects. Our study showed further manipulation of methods may help to improve a clinically acceptable form of drug formulation for such disorders.

## Acknowledgments

Thanks for the help of Golnaz kamalinia and Parvin mahdaviani from the Department of Pharmaceutics, Faculty of Pharmacy, Tehran University of Medical Sciences, Tehran, Iran

## References

[1] A. Omuro, L. M. DeAngelis, Glioblastoma and other malignant gliomas: a clinical review, Jama, Vol. 310, No. 17, pp. 1842–1850, 2013.

[2] M. Kamali, R. Dinarvand, H. Maleki, H. Arzani, P. Mahdaviani, H. Nekounam, M. Adabi, M. Khosravani, Preparation of imatinib base loaded human serum albumin for application in the treatment of glioblastoma, RSC Advances, Vol. 5, No. 76, pp. 62214–62219, 2015.

[3] C.-Y. Ting, C.-H. Fan, H.-L. Liu, C.-Y. Huang, H.-Y. Hsieh, T.-C. Yen, K.-C. Wei, C.-K. Yeh, Concurrent blood–brain barrier opening and local drug delivery using drug-carrying microbubbles and focused ultrasound for brain glioma treatment, Biomaterials, Vol. 33, No. 2, pp. 704–712, 2012.

[4] S. H. Alavizadeh, J. Akhtari, A. Badiee, S. Golmohammadzadeh, M. R. Jaafari, Improved therapeutic activity of HER2 Affibody-targeted cisplatin liposomes in HER2-expressing breast tumor models, Expert opinion on drug delivery, Vol. 13, No. 3, pp. 325–336, 2016.

[5] T. Mainprize, N. Lipsman, Y. Huang, Y. Meng, A. Bethune, S. Ironside, C. Heyn, R. Alkins, M. Trudeau, A. Sahgal, Blood-brain barrier opening in primary brain tumors with non-invasive MR-guided focused ultrasound: a clinical safety and feasibility study, Scientific reports, Vol. 9, No. 1, pp. 1–7, 2019.

[6] A. Parodi, J. Miao, S. M. Soond, M. Rudzinska, A. A. Zamyatnin, Jr., Albumin Nanovectors in Cancer Therapy and Imaging, Biomolecules, Vol. 9, No. 6, pp. 218, Jun 5, 2019.

[7] A. Cseke, T. Schwarz, S. Jain, S. Decker, K. Vogl, E. Urban, G. F. Ecker, Propafenone analogue with additional H-bond acceptor group shows increased inhibitory activity on P-glycoprotein, Archiv der Pharmazie, pp. e1900269, 2020.

[8] L. Jena, E. McErlean, H. McCarthy, Delivery across the blood-brain barrier: nanomedicine for glioblastoma multiforme, Drug delivery and translational research, pp. 1–15, 2019.

[9] H. Nekounam, Z. Allahyari, S. Gholizadeh, E. Mirzaei, M. A. Shokrgozar, R. J. M. S. Faridi-Majidi, E. C, Simple and robust fabrication and characterization of conductive carbonized nanofibers loaded with gold nanoparticles for bone tissue engineering applications, Vol. 117, pp. 111226, 2020.

[10] H. Nekounam, H. Samadian, F. J. b. Asghari, Electro-conductive carbon nanofibers containing ferrous sulfate for bone tissue engineering, 2020.

[11] H. Nekounam, S. Gholizadeh, Z. Allahyari, H. Samadian, N. Nazeri, M. A. Shokrgozar, R. J. M. R. B. Faridi-Majidi, Electroconductive Scaffolds for Tissue Regeneration: Current opportunities, pitfalls, and potential solutions, pp. 111083, 2020.

[12] H. Arzani, M. Adabi, J. Mosafer, F. Dorkoosh, M. Khosravani, H. Maleki, H. Nekounam, M. J. B. R. i. A. C. Kamali, Preparation of curcumin-loaded PLGA nanoparticles and investigation of its cytotoxicity effects on human glioblastoma U87MG cells, Vol. 9, No. 5, pp. 4225–4231, 2019.

[13] Y. Zhang, T. Sun, C. Jiang, Biomacromolecules as carriers in drug delivery and tissue engineering, Acta Pharmaceutica Sinica B, Vol. 8, No. 1, pp. 34–50, 2018.

[14] S. C. Sozer, T. O. Egesoy, M. Basol, G. Cakan-Akdogan, Y. J. J. o. D. D. S. Akdogan, Technology, A simple desolvation method for production of cationic albumin nanoparticles with improved drug loading and cell uptake, Vol. 60, pp. 101931, 2020.

[15] S. Abbasi, A. Paul, W. Shao, S. J. J. o. d. d. Prakash, Cationic albumin nanoparticles for enhanced drug delivery to treat breast cancer: preparation and in vitro assessment, Vol. 2012, 2012.

[16] B. Kim, C. Lee, E. S. Lee, B. S. Shin, Y. S. Youn, Paclitaxel and curcumin co-bound albumin nanoparticles having antitumor potential to pancreatic cancer, asian journal of pharmaceutical sciences, Vol. 11, No. 6, pp. 708–714, 2016.

[17] V. J. Muniswamy, N. Raval, P. Gondaliya, V. Tambe, K. Kalia, R. K. Tekade, ‘Dendrimer-Cationized-Albumin’encrusted polymeric nanoparticle improves BBB penetration and anticancer activity of doxorubicin, International journal of pharmaceutics, Vol. 555, pp. 77–99, 2019.

[18] S. A. Y. Rehana H, Iqbal QA, Kalsoom R, khalid Iqbal R, Anwar F, A Review: Targeted Cancer Therapy as a Fight Against Brain Tumor, American Journal of Biomedical Science and Research, Vol. 3, pp. 5, 2019.

[19] Y. Liu, W. Lu, Recent advances in brain tumor-targeted nano-drug delivery systems, Expert opinion on drug delivery, Vol. 9, No. 6, pp. 671–686, 2012.

[20] E. N. Hoogenboezem, C. L. Duvall, Harnessing albumin as a carrier for cancer therapies, Advanced drug delivery reviews, Vol. 130, pp. 73–89, 2018.

[21] L. Van de Sande, S. Cosyns, W. Willaert, W. Ceelen, Albumin-based cancer therapeutics for intraperitoneal drug delivery: a review, Drug Delivery, Vol. 27, No. 1, pp. 40–53, 2020.

[22] W. M. Pardridge, New approaches to drug delivery through the blood-brain barrier, Trends in biotechnology, Vol. 12, No. 6, pp. 239–245, 1994.

[23] V. J. Muniswamy, N. Raval, P. Gondaliya, V. Tambe, K. Kalia, R. K. J. I. j. o. p. Tekade, ‘Dendrimer-Cationized-Albumin’encrusted polymeric nanoparticle improves BBB penetration and anticancer activity of doxorubicin, Vol. 555, pp. 77–99, 2019.

[24] M. K. Riley, W. Vermerris, Recent advances in nanomaterials for gene delivery—a review, Nanomaterials, Vol. 7, No. 5, pp. 94, 2017.

[25] A. Parodi, M. Rudzińska, A. A. Deviatkin, S. M. Soond, A. V. Baldin, A. A. Zamyatnin, Established and emerging strategies for drug delivery across the blood-brain barrier in brain cancer, Pharmaceutics, Vol. 11, No. 5, pp. 245, 2019.

[26] W. Lohcharoenkal, L. Wang, Y. C. Chen, Y. Rojanasakul, Protein nanoparticles as drug delivery carriers for cancer therapy, BioMed research international, Vol. 2014, 2014.

[27] A. M. Diels, C. W. J. C. r. i. m. Michiels, High-pressure homogenization as a non-thermal technique for the inactivation of microorganisms, Vol. 32, No. 4, pp. 201–216, 2006.

[28] S. Schultz, G. Wagner, K. Urban, J. J. C. E. Ulrich, T. I. C. P. E. P. Engineering-Biotechnology, High-pressure homogenization as a process for emulsion formation, Vol. 27, No. 4, pp. 361–368, 2004.

[29] S. M. Jafari, E. Assadpoor, Y. He, B. J. F. h. Bhandari, Re-coalescence of emulsion droplets during high-energy emulsification, Vol. 22, No. 7, pp. 1191–1202, 2008.

[30] K. S. Yadav, K. J. J. o. P. I. Kale, High pressure homogenizer in pharmaceuticals: understanding its critical processing parameters and applications, pp. 1–12, 2019.

[31] H. L. Tan, L. V. J. J. o. M. L. Woodcock, Molecular dynamics study of a simple liquid at negative pressures, Vol. 136, No. 3, pp. 281–287, 2007.

[32] J. Jeevanandam, Y. San Chan, M. K. J. B. Danquah, Nano-formulations of drugs: recent developments, impact and challenges, Vol. 128, pp. 99–112, 2016.

[33] M. G. Dilshara, I. M. N. Molagoda, R. G. P. T. Jayasooriya, Y. H. Choi, C. Park, K. T. Lee, S. Lee, G.-Y. Kim, p53-Mediated Oxidative Stress Enhances Indirubin-3′-Monoxime-Induced Apoptosis in HCT116 Colon Cancer Cells by Upregulating Death Receptor 5 and TNF-Related Apoptosis-Inducing Ligand Expression, Antioxidants, Vol. 8, No. 10, pp. 423, 2019.

[34] K. Misumi, T. Ogo, J. Ueda, A. Tsuji, S. Fukui, N. Konagai, R. Asano, S. Yasuda, Development of pulmonary arterial hypertension in a patient treated with Qing-Dai (Chinese herbal medicine), Internal Medicine, Vol. 58, No. 3, pp. 395–399, 2019.

[35] L. Chen, J. Wang, J. Wu, Q. Zheng, J. Hu, Indirubin suppresses ovarian cancer cell viabilities through the STAT3 signaling pathway, Drug design, development and therapy, Vol. 12, pp. 3335, 2018.

[36] A. Rahiminejad, R. Dinarvand, B. Johari, S. J. Nodooshan, A. Rashti, E. Rismani, P. Mahdaviani, Z. Saltanatpour, S. Rahiminejad, M. Raigani, Preparation and investigation of indirubin-loaded SLN nanoparticles and their anticancer effects on human glioblastoma U87MG cells, Cell biology international, Vol. 43, No. 1, pp. 2–11, 2019.

[37] M. Thöle, S. Nobmann, J. Huwyler, A. Bartmann, G. Fricker, Uptake of cationized albumin coupled liposomes by cultured porcine brain microvessel endothelial cells and intact brain capillaries, Journal of drug targeting, Vol. 10, No. 4, pp. 337–344, 2002.

[38] H. Lu, L. Noorani, Y. Jiang, A. W. Du, M. H. Stenzel, Penetration and drug delivery of albumin nanoparticles into pancreatic multicellular tumor spheroids, Journal of Materials Chemistry B, Vol. 5, No. 48, pp. 9591–9599, 2017.

[39] N. P. Desai, C. Tao, A. Yang, L. Louie, T. Zheng, Z. Yao, P. Soon-Shiong, S. Magdassi, Protein stabilized pharmacologically active agents, methods for the preparation thereof and methods for the use thereof, Google Patents, 1999.

[40] B. Chertok, B. A. Moffat, A. E. David, F. Yu, C. Bergemann, B. D. Ross, V. C. Yang, Iron oxide nanoparticles as a drug delivery vehicle for MRI monitored magnetic targeting of brain tumors, Biomaterials, Vol. 29, No. 4, pp. 487–496, 2008.

[41] H. J. Byeon, S. Lee, S. Y. Min, E. S. Lee, B. S. Shin, H.-G. Choi, Y. S. Youn, Doxorubicin-loaded nanoparticles consisted of cationic-and mannose-modified-albumins for dual-targeting in brain tumors, Journal of Controlled Release, Vol. 225, pp. 301–313, 2016.

[42] W. Lu, Y. Zhang, Y.-Z. Tan, K.-L. Hu, X.-G. Jiang, S.-K. Fu, Cationic albumin-conjugated pegylated nanoparticles as novel drug carrier for brain delivery, Journal of controlled release, Vol. 107, No. 3, pp. 428–448, 2005.

[43] S. Das, R. Banerjee, J. Bellare, Aspirin loaded albumin nanoparticles by coacervation: implications in drug delivery, Trends Biomater Artif Organs, Vol. 18, No. 2, pp. 203–12, 2005.

[44] M. Hua, X. Hua, Polymer nanoparticles prepared by supercritical carbon dioxide for in vivo anti-cancer drug delivery, Nano-Micro Letters, Vol. 6, No. 1, pp. 20–23, 2014.

